# The effects of development and chronic oxidative stress on telomere length in an agricultural pest moth, *Helicoverpa armigera*

**DOI:** 10.1101/2020.12.15.422848

**Authors:** Nonthakorn (Beatrice) Apirajkamol, Tom K Walsh, Angela McGaughran

## Abstract

Telomeres are repetitive sequences located at the end of chromosomes in eukaryotes that protect against loss of important sequences during the cell replication process. Telomere length (TL) shortens with every round of cell division. When a telomere becomes too short, cells can no longer proliferate and this triggers the cell apoptosis process. Apart from cell replication, the length of telomeres can be affected by factors such as sex, genetics, and stress levels. Oxidative stress in particular can cause damage to telomeres and telomere maintenance processes, resulting in TL shortening. This phenomenon occurs in humans and many vertebrates, especially endothermic species. However, the ways in which various stress types affect the TL of invertebrate species remains ambiguous.

Here, we examined the effects of development and oxidative stress on TL in the invertebrate pest moth, *Helicoverpa armigera*. In the former case, we extracted genomic DNA from three developmental stages (1-day old egg, 4th instar, and first-day emerged moths) and measured TL by qPCR. In the latter, we chronically exposed individuals to paraquat – an organic herbicide that induces oxidative stress - and then measured TL as per our development methodology. In addition, we examined TL in a subset of published whole genome short-read sequencing data of caterpillars and moths using the software, Computel.

In our experimental work, we found that TL in *H. armigera* was significantly longer at the early stages of development and shortens in later stages. However, oxidative stress does not appear to shorten TL in *H. armigera* following chronic exposure to paraquat. In our Computel analysis, we found that caterpillars had longer mean TL than moths but this difference was not significant due to the high variation among samples.

Collectively, our research provides new data on TL in an underrepresented group, adding new insights into the progression of TL shortening with development and the effects of oxidative stress on TL, while also more generally highlighting the value of applying complementary approaches to TL measurement.

## Introduction

Telomeres are repetitive non-coding sequences that protect chromosomes from losing functional regions. Eukaryotes have linear chromosomes that cannot be synthesised at the very ends during the cell division process. Therefore, chromosomes lose small amounts of DNA sequence with every round of cell replication [1]. The majority of eukaryotes can partially compensate for this with an enzyme called telomerase, which can synthesise the repeat sequence [2]. Telomerase is ordinarily highly expressed during developmental periods and in certain types of cells (e.g., reproductive and malignant cells [3]); however, the activity of telomerase is negligible in stem cells and absent in normal adult somatic cells [4]. As a result, in many species, telomere length (TL) shortens over time [1]. When telomeres eventually become too short to replicate, the cell will stop proliferating and enter senescence or apoptosis [5] and this is part of the process leading to aging. Thus, shortening of telomeres is associated with several age-related diseases in humans [6, 7, 8], and also shortens the lifespan of some species (e.g., [9, 10]).

TL is not only dependant on the number of replication/cell division cycles, but is also affected by genetic factors, sex, and other external stimuli, such as chronic stress [11]. In particular, there is some evidence to support a correlation between chronic stress and shortened telomeres in humans [12]. For example, people who have suffered from depression or anxiety may have shorter TLs [13, 14], and childhood trauma [15], bipolar disorder [16], physical and sexual assault [17], and PTSD (Post-Traumatic Stress Disorder; [18]) have also been shown to affect TL.

Although the mechanism remains unclear, in many organisms, stress appears to affect the length of telomeres by lowering telomerase activity or enhancing oxidative stress. In humans, early exposure in life to chronic stress may affect TL by down-regulating telomerase expression to reduce telomerase activity [19, 20, 21]. This may account for the apparent link between stress and shorter lifespans i.e., individuals experiencing stress during early development have less telomerase activity and therefore shorter TL leading to a reduced lifespan. Chronic stress can also generate reactive oxygen species (ROS), resulting in a disruption of the antioxidant/oxidant balance and a subsequent increase in oxidative stress [22]. High levels of ROS can affect the activity of telomerase by oxidising the guanine in telomeres, preventing the telomerase from extending telomeres during development [23, 24] and ROS are also more likely to cleave repeated telomeric sequences than non-telomeric regions [25].

The relationship between TL and stress in humans has been studied for more than a decade and indications are that stress can have long-term effects on human TL that can even be passed to offspring [19, 26, 27]. However, recent research has shown that the relationship is highly complex and context dependent, with the generality of stress causing TL attrition being questioned, alongside research that shows that associations can be transient (e.g., [28]), can strongly depend on the tissue-type, life-stage, and experimental condition being tested [29, 30], and may be affected by the method of telomere measurement [31]. It is even less clear how these processes might work in other species, with telomere dynamics and maintenance systems in invertebrate and ectothermic species in particular varying or remaining unclear due to a lack of research.

*Helicoverpa armigera* is a pest moth that is widespread across several continents [32]. The larvae of this species damages the reproductive part of crops, leading to significant economic damage annually [33]. However, prevention and management of *H. armigera* infestation is a challenge due to their wide host range, high fecundity, and ability to migrate over long distances and evolve resistance to insecticides [33, 34, 35]. Here, we aim to reveal the effects of stress on TL in *H. armigera* following chronic exposure to paraquat (an oxidative stress producer). Paraquat (*N,N*′-dimethyl-4,4′-bipyridinium dichloride) is an organic herbicide that exerts its toxic effect by generating super oxide anions [36]. Despite being banned in several countries, paraquat is widely available throughout the world due to its efficiency and low cost [37]. As a result, paraquat is found on several plant species and may be present in the natural diet of *H. armigera* in certain parts of the world.

Oxidative stress has been shown to shorten TL in mice [38]. In addition, there is evidence that paraquat in particular has a strong impact on development, gene expression, and TL in insects. For example, in *D. melanogaster*, paraquat has been shown to increase mortality, reduce climbing ability, and result in up-regulation of several antioxidant genes [39], as well as reduce TL elongation rates [40]. In *H. armigera*, paraquat injected into pupae has been shown to extend diapause by affecting the insulin-signalling pathway [41], while our own previous research demonstrated that chronic oxidative stress exposure delays development and alters gene expression in this species [42]. Here, we use *H. armigera* as a model system to further investigate chronic oxidative stress by examining its effects on TL.

## Materials and Methods

### Study samples and oxidative stress exposure

Samples in this study were obtained from a laboratory colony at the Commonwealth Scientific and Industrial Research Organisation (CSIRO) in Canberra, Australia. This colony was origianlly isolated from Northern New South Wales, Australia. *H. armigera* were exposed to paraquat, leading to chronic oxidative stress (described in detail in [42]). In brief, experiments used 2-day old fertilised eggs collected from 40 healthy moths, air-dried, and allowed to hatch. Caterpillars (1^st^-instar) were reared until pupation under optimal laboratory conditions (25 ± 1°C, 50 ± 10% relative humidity, and light day:night 14:10 to imitate natural light) on artificial diet, which was changed weekly. Moths were fed a honey solution. Based on previous work [42], paraquat concentrations of 0.3 and 0.4 mM were chosen for our ‘stressed’ samples, which were reared as outlined above for controls and chronically exposed to paraquat through their diet (i.e., incorporation of the required concentration into the solid artificial diet before it set, or directly into the honey solution) upon hatching.

### DNA extraction

To compare TL between different developmental stages and treatments, six individuals for three developmental stages (1-day old egg, 4^th^ instar, and newly emerged moths) from control and stressed samples were collected in absolute ethanol and stored at −20°C. Second generation eggs were also collected from both controls and stressed moths to assess the spill over effect of stress on TL in the next generation.

In order to isolate genomic DNA (gDNA), samples were air-dried, allowing ethanol to evaporate at room temperature. Because TL might shorten differently in different tissue types, the midgut of 4^th^-instar caterpillars was isolated, as this tissue has high contact with paraquat after ingestion. Leg tissue of moths was chosen to avoid the reproductive system in adults. Dried tissue was placed into 1.5 mL Eppendorf tubes. Lysis buffer was added and samples were ground with DNase-free plastic pestles before 20 μl of proteinase K was added and samples were incubated overnight at 58°C. DNA extraction was then conducted following the JetFlex™Genomic DNA Purification Kit DNA (Cat no. A30701) protocol to isolate gDNA from tissue samples. Subsequently, gDNA was re-suspended by adding 20 μl (eggs and moth legs) or 50 μl (4^th^-instar midguts) of TE buffer and incubating overnight at room temperature before storage at −20°C.

### Telomere length measurement via qPCR

TL in both control and stressed samples (six biological replicates each) for both the current (eggs, 4^th^-instar, moths) and 2^nd^ generation (eggs) was determined by qPCR following the Luna^®^ Universal qPCR Master Mix Protocol (M3003, New England Biolabs) using 2 μl of DNA template. The primers were designed based on telomeric sequence of other Lepidopteran species (TTAGG)_n_ [43, 44, 45]. This sequence was also confirmed by mining it from existing long-read sequencing (minION) data of *H. armigera* obtained from the Australian National Insect Collection (ANIC; Dr. Luana S. F. Lins). For comparison, elongation factor 1-alpha (EF1, GenBank accession No. U20129) was chosen as a housekeeping gene not expected to have changed in length over time or treatment. The primers used were Tel F: GGTTTTTTGGGTTGGGTTGGGTTGGGTTGGGTT, Tel R: GGCTTGCCTTACCTTACCTTACCTTACCTTACCT, EF1 F: GACAAACGTACCATCGAGAAG, and EF1 R: GATACCAGCCTCGAAGTCAC.

Mean T/S ratio (i.e., the ratio of the telomere repeat copy number to the single-copy - elongation factor 1-alpha – number) was calculated following Livak’s equation [46, 47] for each developmental stage and treatment.

### Telomere length computation with Computel

The software, Computel ver. 1.2 (available at: https://github.com/lilit-nersisyan/computel; [48]) was used to compute mean telomere length from whole genome re-sequencing data for 50 individuals of *H. armigera* collected from ten countries around the world (Australia, Brazil, China, France, India, Madagascar, New Zealand, Senegal, Spain, Uganda), as larvae from wild and crop host plants or as adult moths via light/pheromone traps [49]. This included 23 caterpillars and 27 moths, although we lack information as to the ages of any of the samples beyond this broad categorisation. The software was run in default mode on paired-end fastq files obtained from [49], with the number of chromosomes, genome length, and telomere repeat pattern specified.

### Statistical analysis

In order to evaluate differences in TL between developmental stages and between control and stressed samples, results were statistically analysed in SPSS ver. 22 [50]. Means and standard deviations (SD) were calculated for all measures and one-way ANOVA with Tukey’s Honestly Significant Difference (HSD) was used to assess differences between groups at a 95% confidence interval. In this analysis, samples that are statistically similar (in terms of mean and variance for TL) group together into homogeneous subsets. Thus, if samples are categorised into different groups (referred to as ‘a’ or ‘b’ see Results), they are considered significantly different from each other. Data visualisation was performed using the ggplot2 ver. 3.2.1 [51] package in R ver 3.6.1 [52].

To evaluate differences in mean TL between caterpillars and moths, as computed in Computel, an unpaired t-test was performed in R. A Pearson’s product-moment correlation test was also performed using the cor.test command in R to examine whether estimates of telomere length were associated with base coverage [48]. Finally, mean computed telomere length per sample, and the correlation between mean telomere length and base coverage, were plotted with ggplot2 in R.

## Results

### Telomere length during development

Differences in the amount of relative telomeric sequence (i.e. T/S ratio) of *H. armigera* at different developmental stages and across experimental treatments are illustrated in Figure 1.

**Figure 1.**
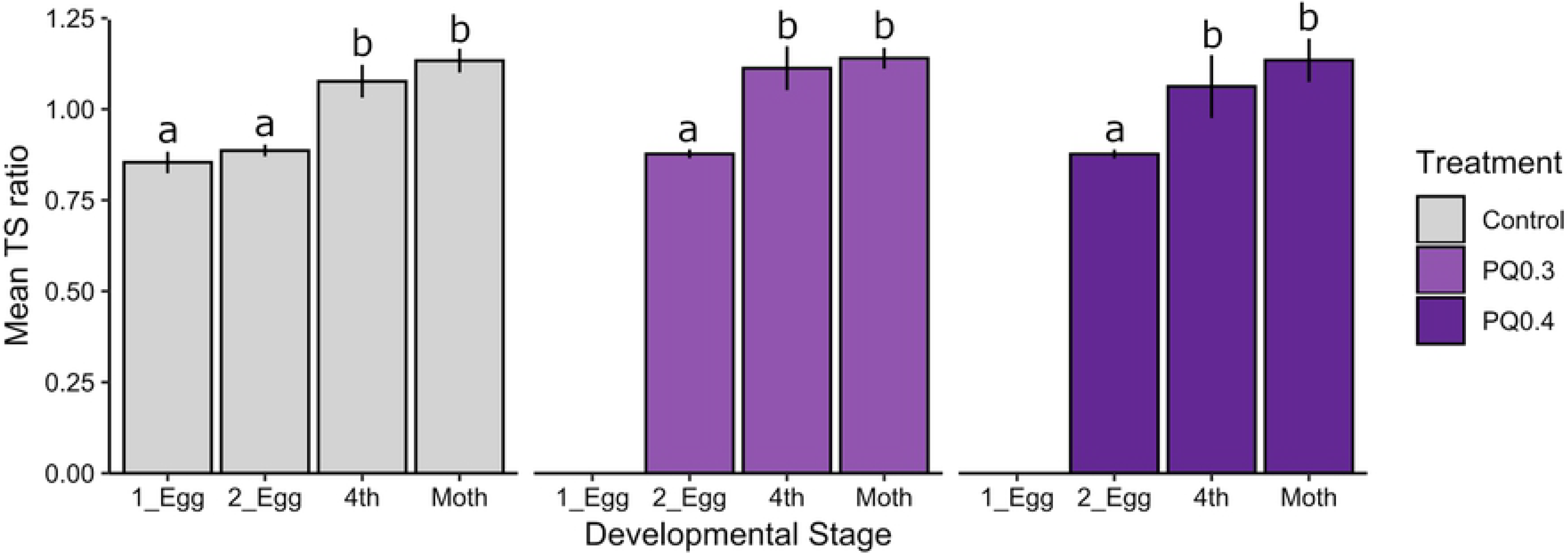
Effects of development and stress on telomere length in *Helicoverpa armigera*. Mean TS ratio (i.e., the ratio of the telomere repeat copy number to the single-copy - elongation factor 1-alpha – number) following chronic oxidative stress exposure for developmental stage and treatment as shown by the *x-axis* and legend, respectively. Labels on the *x-axis* correspond to: ‘1_Egg’ = 1^st^ generation eggs; ‘2_Egg’ = 2^nd^ generation eggs; 4^th^ = 4^th^-instar. PQ0.3 and PQ0.4 correspond to use of paraquat doses of 0.3 mM and 0.4 mM, respectively. Significant differences among developmental stages/treatments are indicated by non-overlapping characters (‘a’, ‘b’), and error bars indicate standard deviation.

The relative TL of whole eggs (‘1_Egg’) and other developmental stages (‘4^th^’ and ‘Moth’) were significantly different (as indicated by ‘a’ and ‘b’ subgroups in the figure), with TL in 4^th^-instar individuals shorter than TL of eggs for both control and stressed samples (F_8,49_=38.679, *P* < 0.001). However, 4^th^-instar and moth TLs of controls were not significantly different from each other (both in ‘b’ subset).

These results were consistent with the results of the Computel analysis, where mean TL was found to be longer in caterpillars compared to moths, though there was a lot of variability in mean TL among samples and this result was not significant (mean of caterpillar minus moth = 959.000; 95% confidence interval of difference: −683.608:2601.608; T_48_ = 1.175; *P* = 0.246; SE of difference = 816.462) (Fig. 2A). A correlation analysis found that mean telomere length was significantly associated with base coverage per sample (T_48_ = −2.330; *P* = 0.024; 95% confidence interval: −0.548:−0.044; R = −0.319), such that lower estimates of base coverage resulted in higher estimates of telomere length (Fig. 2B).

**Figure 2.**
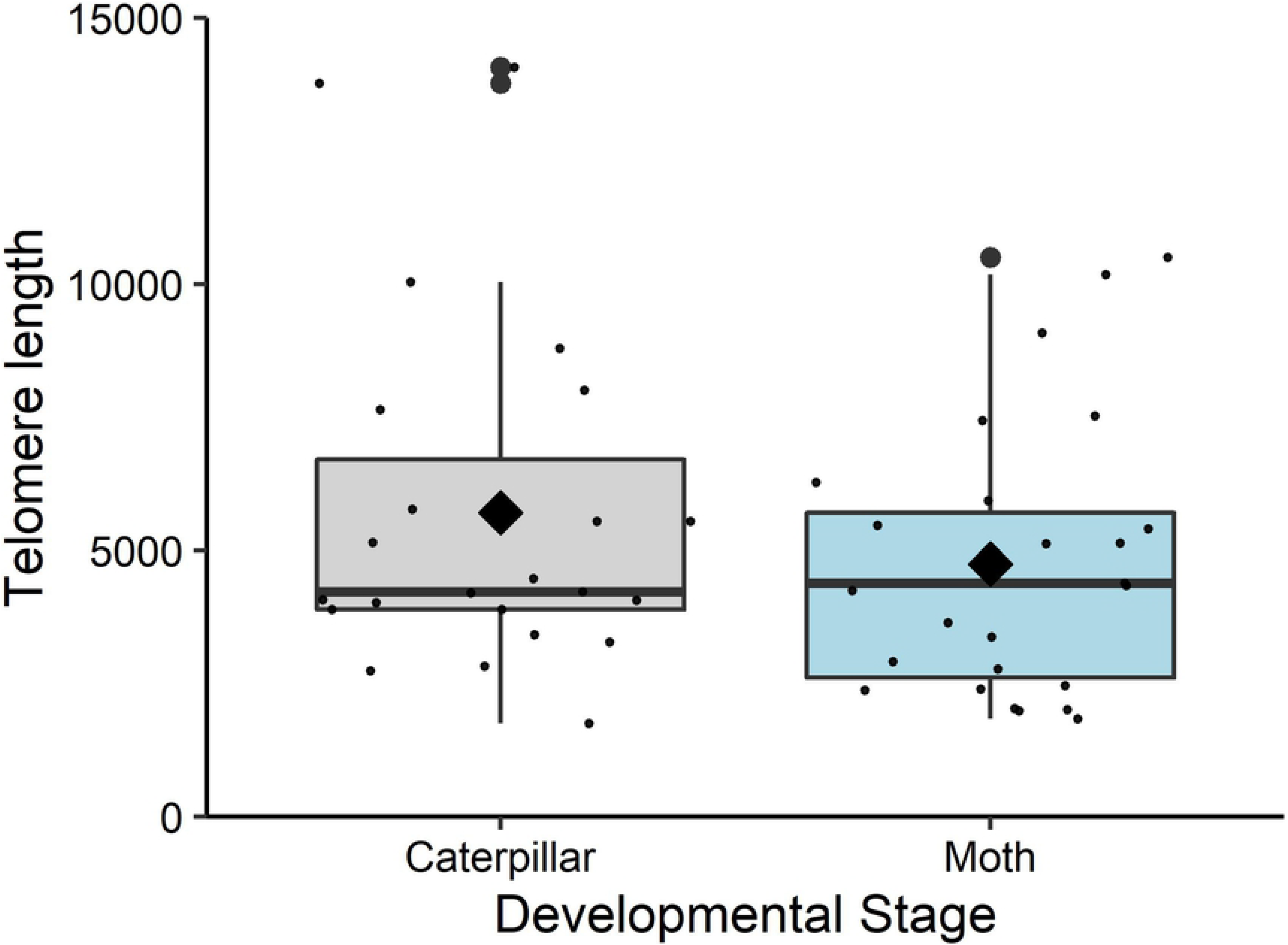
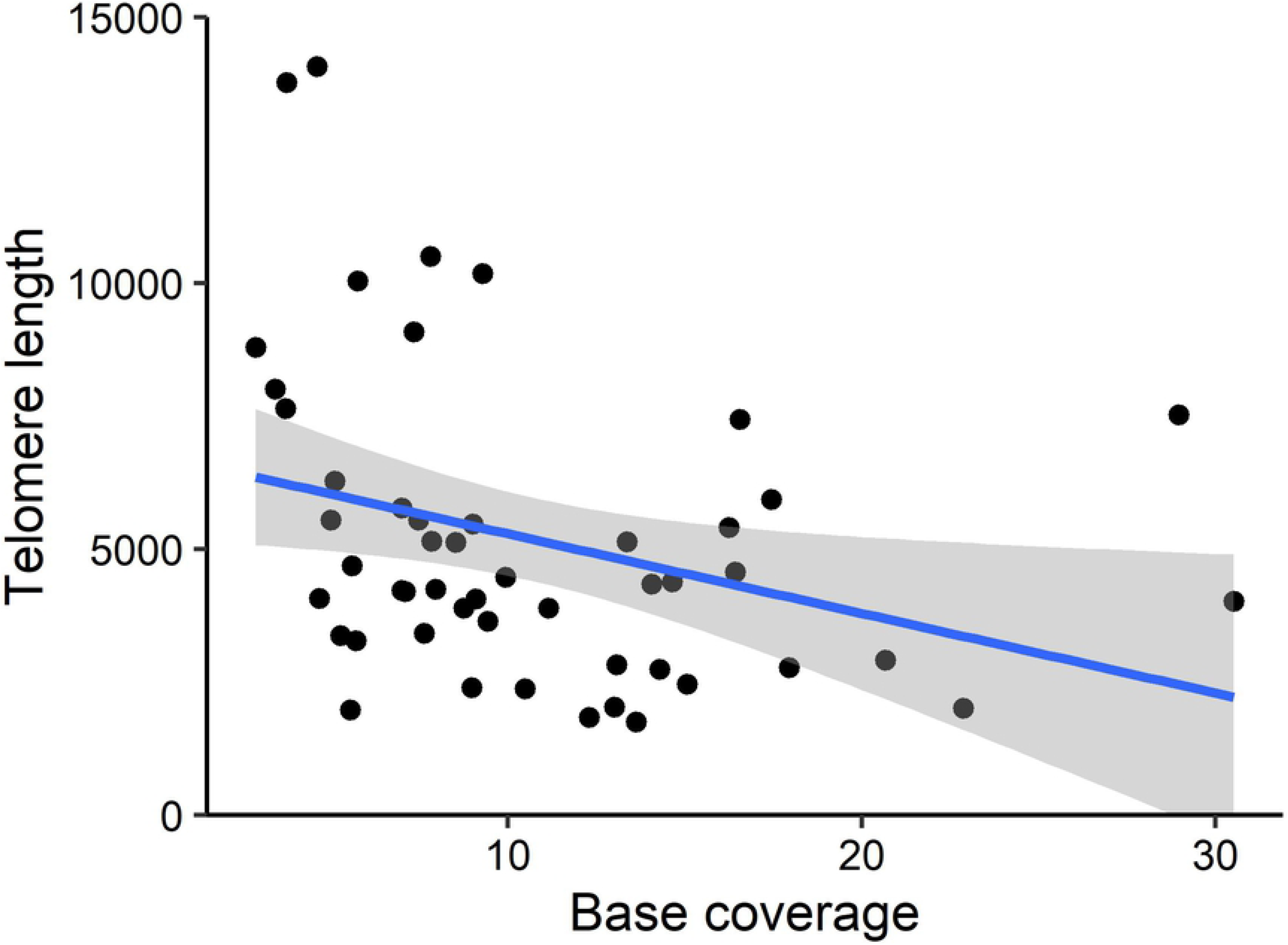
Computed mean telomere length in *Helicoverpa armigera*. Whole genome re-sequencing data was processed in the software, Computel, to compute mean telomere length for samples collected from the field and broadly classified as ‘caterpillar’ and ‘moth’. (A) The boxplots show the distribution of mean telomere lengths across samples for each of these categories (with the mean indicated by a black diamond inside the boxplot), indicating a general but non-significant (T_48_ = 1.175; *P* = 0.246), trend of longer telomeres in caterpillars compared to moths; (B) The correlation between mean telomere length and base coverage per sample (T_48_ = −2.330; *P* = 0.024; 95% confidence interval: −0.548:−0.044; R = −0.319). Raw data points are plotted, along with the linear regression trend line.

### Telomere length in response to oxidative stress

The relative TLs of control and paraquat 0.3 mM and 0.4 mM stressed samples were not significantly different for the 4^th^-instar and moth stages (all ‘b’ subset in Fig. 1), or for the eggs of the next generation (‘2_Egg’; all ‘a’ subset in Fig. 1) (F_3,20_=2.645, *P* = 0.077).

## Discussion

There is a large body of work that suggests that stress can have a negative effect on TL in various species. However, our results suggest that, while aging negatively correlates with TL in *H. armigera*, chronic oxidative stress does not, at least under the experimental conditions we applied.

In particular, we found that TL is dramatically shorter in 4^th^-instar individuals compared to eggs, though not between 4th-instar and moths. This finding may relate to the caterpillar stage being when *H. armigera* undergoes significant growth (see [42]), and therefore cell division and proliferation compared to the other developmental stages. As a pupa and adult, there is also less nutritional uptake (pupae are non-feeding and adults consume only small amounts of honey solution), leading to less exosure to ROS production and its associated effects on telomerase inhibition and telomere damage [23, 24].

This experimental result was consistent with our computation of mean TLs from whole genome re-sequencing data, which generally indicated that mean TL is shorter in moths compared to caterpillars. Although this trend was not significant for our data, there was a lot of variation in the dataset and we lack information as to the specific ages of the tested individuals beyond their broad categorisation as ‘caterpillar’ and ‘moth’ as they were all field-collected vs. controlled for development under laboratory conditions. In addition, the estimates of mean telomere length were significantly correlated with estimates of per sample base coverage. In general, increasing coverage is known to improve the accuracy of mean TL estimation in Computel, however performance comparisons with synthetic data have shown that the accuracy of Computel was not affected for 100 bp read lengths and coverage values of 0.2x, 2x and 10x [48] – our read length was 90 or 101, and our base coverage ranged from 3x to 31x. Overall, both our experimental and computational results are consistent with the hypothesis that TL shortens as a result of aging, as has been shown in other species, including black garden ants [53], humans [1], and birds [54].

Conversely, TL was not found to be shortened in response to chronic oxidative stress in the current study and we saw no spillover effects in second generation eggs. According to the literature, various types of stress can shorten TL in a variety of species, including humans [55, 56, 57], rodents [58, 38], birds [59, 54, 60, 61, 62, 63], fish [64, 65], lizards [66], toads [67], flies [40], plants [68], and yeast [69, 70]. However, other studies have found no such correlation (nine-spined stickleback, [71]; young brown trout, [72]; desert agama lizard, [66], Western spadefoot toad, [67]; garden snails, [73]), or even elongated TL with various stress types (e.g., in yeast; [69]).

Although the reason for the lack of relationship in the fish remains unclear, for yeast and the desert lizard, the authors each suggested that effects of stress on TL may depend on previous exposure to the particular stress applied. For example, [69] found that *S. cerevisiae* experienced TL elongation under alcohol stress, and [66]) found that consistent thermal stress in the desert lizard led to an up-regulated thermal-regulatory ability to protect TL integrity. Because alcohol (yeast) and consistent thermal stress (desert lizard) are regularly faced by these species in their natural environments, they may have mechanisms in place to quickly mitigate these stresses. Similarly, our finding of no significant differences in TL between control and stressed samples of *H. armigera* may be due to the fact that oxidative stress is induced through the plant defence mechanisms that *H. armigera* naturally faces in the wild [74]. Thus, *H. armigera* may have a well-developed and effective mechanism to cope with chronic oxidative stress. It is unclear what such a mechanism may be, however in our previous work using the same experimental samples to examine gene expression, we found that telomerase genes were not up-regulated in paraquat stressed samples [42]. It is possible that acute severe stress, such as pesticide exposure, would lead to shortening of telomeres in *H. armigera*, but this remains to be investigated.

There is some also evidence to suggest that different levels of stress may affect TL differently in some species. For example, [40] stressed *Drosophila melanogaster* with paraquat at different concentrations and then measured the telomeric retroelements (NB: *Drosophila* has a different telomeric system to *H. armigera*; [75]). These authors found that only certain non-/sub-lethal concentrations of paraquat increased TL in *Drosophila*, whereas too much or too little paraquat resulted in no significant effects. Interestingly, some TL shortening effects in *D. melanogaster* were also shown in this study to occur only after multi-generational exposure to paraquat, suggesting that the effects of such stress on TL might be too minor to determine in a single generation [40]. This is an exciting result that could be explored with future work.

Thus, the lack of a relationship between chronic oxidative stress and TL found here may be due to the chosen experimental design (type or concentration of stress applied). It is also possible that the effects of such stress on TL in *H. armigera* are very minor or transient and that the TL measurement approaches used in this experiment were not sensitive enough to detect them. However, because the methods applied did see the expected decrease in TL with age, we believe this latter explanation to be less likely. More generally, as noted in the Introduction, the relationship between stress and TL is known to be highly complex and context (tissue-type, life-stage, stress-type, TL measurement method, etc.) dependent [29, 31, 30].

In summary, we provide new evidence of TL shortening with development in *H. armigera* that is consistent with the general hypothesis that aging is associated with TL attrition. Chronic oxidative stress, however, does not appear to shorten TL in *H. armigera* in the current or second generation under the conditions of chronic paraquat exposure used here. This is consistent with recent research that calls the generality of TL attrition with stress into question and highlights the context-dependent nature of TL dynamics.

## Acknowledgements

We would like to thank Kerensa McElroy for early discussions about calculating TL from whole genome resequencing data; Bill James for assistance with rearing of *Helicoverpa armigera*; John Roberts, Egi Kardia and David Heckel for suggestions regarding TL measurement; Luana Lins for confirming the telomeric sequence of *H. armigera* from minION data; and Amanda Padovan, Rachael Remington, Theodore Colls, and the NBA’s thesis advisory panel (Rod Peakall, Maja Adamska, Benjamin Schwessinger) for feedback on earlier drafts of this work. This project was supported through funding from the Australian Research Council (Discovery Early Career Researcher Award DE160100685 to AM), the Centre for Biodiversity Analysis (Ignition Grant to AM), and the Commonwealth Scientific and Industrial Research Organisation (Land and Water).

## References

[1] Blackburn EH. Telomere states and cell fates. Nature. 2000;408: 53–56.

[2] Chan SR, Blackburn EH. Telomeres and telomerase. Philosophical Transactions of the Roy Soc London B: Biolog Sci. 2004;359: 109–122.

[3] Holt SE, Wright WE, Shay JW. Regulation of telomerase activity in immortal cell lines. Mol Cell Biol. 1996;16: 2932–2939.

[4] Weng NP, Levine BL, June CH, Hodes RJ. Regulated expression of telomerase activity in human T lymphocyte development and activation. J Exp Med. 1996;183: 2471–2479.

[5] Szostak JW, Blackburn EH. Cloning yeast telomeres on linear plasmid vectors. Cell. 1982;29: 245–255.

[6] Brouilette S, Singh RK, Thompson JR, Goodall AH, Samani NJ. White cell telomere length and risk of premature myocardial infarction. Arterioscler Thromb Vasc Biol. 2003;23: 842–846.

[7] Hathcock KS, Jeffrey Chiang Y, Hodes RJ. In vivo regulation of telomerase activity and telomere length. Immunolog Rev. 2005;205: 104–113.

[8] Aviv A, Valdes A, Gardner JP, Swaminathan R, Kimura M, Spector TD. Menopause modifies the association of leukocyte telomere length with insulin resistance and inflammation. J Clin Endo Metab. 2006;91: 635–640.

[9] Haussmann MF, Mauck RA. Telomeres and longevity: testing an evolutionary hypothesis. Mol Biol Evol. 2008;25: 220–228.

[10] Bize P, Criscuolo F, Metcalfe NB, Nasir L, Monaghan P. Telomere dynamics rather than age predict life expectancy in the wild. Proc Roy Soc B: Biolog Sci. 2009;276: 1679–1683.

[11] Spivak IM, Mikhelson VM, Spivak DL. Telomere length, telomerase activity, stress, and aging. Adv Gerontol. 2016;6: 29–35.

[12] Mitchell C, Hobcraft J, McLanahan SS, Siegel SR, Berg A, Brooks-Gunn J, et al. Social disadvantage, genetic sensitivity, and children’s telomere length. Proc Natl Acad Sci. 2014;111: 5944–5949.

[13] Cai N, Chang S, Li Y, Li Q, Hu J, Liang J, et al. Molecular signatures of major depression. Curr Biol. 2015;25: 1146–1156.

[14] Li J, Hou R, Niu X, Liu R, Wang Q, Wang C, et al. Comparison of microarray and RNA-Seq analysis of mRNA expression in dermal mesenchymal stem cells. Biotechnol Lett. 2016;38: 33–41.

[15] Tyrka AR, Price LH, Kao HT, Porton B, Marsella SA, Carpenter LL. Childhood maltreatment and telomere shortening: preliminary support for an effect of early stress on cellular aging. Biolog Psych. 2010;67: 531–534.

[16] Lima IM, Barros A, Rosa DV, Albuquerque M, Malloy-Diniz L, Neves FS, et al. Analysis of telomere attrition in bipolar disorder. J Affect Dis. 2015;172: 43–47.

[17] Shalev I, Entringer S, Wadhwa PD, Wolkowitz OM, Puterman E, Lin J, et al. Stress and telomere biology: a lifespan perspective. Psychoneuroendocrinology. 2013;38: 1835–1842.

[18] Lohr JB, Palmer BW, Eidt CA, Aailaboyina S, Mausbach BT, Wolkowitz OM, et al. Is post-traumatic stress disorder associated with premature senescence? A review of the literature. Am J Geriatr Psychiatry. 2015;23: 709–725.

[19] Epel ES, Blackburn EH, Lin J, Dhabhar FS, Adler NE, Morrow JD, et al. Accelerated telomere shortening in response to life stress. Proc Natl Acad Sci. 2004;101: 17312–17315.

[20] Damjanovic AK, Yang Y, Glaser R, Kiecolt-Glaser JK, Nguyen H, Laskowski B, et al. Accelerated telomere erosion is associated with a declining immune function of caregivers of Alzheimer’s disease patients. J Immunol. 2007;179: 4249–4254.

[21] Zhou QG, Hu Y, Wu DL, Zhu LJ, Chen C, Jin X, et al. Hippocampal telomerase is involved in the modulation of depressive behaviors. J Neuroscience. 2011;31: 12258–12269.

[22] Mittler R. Oxidative stress, antioxidants and stress tolerance. Trends Plant Sci. 2002;7: 405–410.

[23] Houben JM, Moonen HJ, van Schooten FJ, Hageman GJ. Telomere length assessment: biomarker of chronic oxidative stress? Free Radic Biol Med. 2008;44: 235–246.

[24] Smith S. Telomerase can’t handle the stress. Genes Dev. 2018;32: 597–599.

[25] Von Zglinicki T. Oxidative stress shortens telomeres. Trends Biochem Sci. 2002;27: 339–344.

[26] Gladych M, Wojtyla A, Rubis B. Human telomerase expression regulation. Biochem Cell Biol. 2011;89: 359–376.

[27] Coimbra BM, Carvalho CM, Moretti PN, Mello MF, Belangero SI. Stress-related telomere length in children: a systematic review. J Psych Res. 2017;92: 47–54.

[28] Heidinger BJ, Blount JD, Boner W, Griffiths K, Metcalfe NB, Monaghan P. Telomere length in early life predicts lifespan. Proc Natl Acad Sci. 2012;109: 1743–1748.

[29] Prowse KR, Greider CW. Developmental and tissue-specific regulation of mouse telomerase and telomere length. Proc Natl Acad Sci. 1995;92: 4818–4822.

[30] Rollings N, Friesen CR, Whittington CM, Johansson R, Shine R, Olsson M. Sex-And tissue-specific differences in telomere length in a reptile. Ecol Evol. 2019;9: 6211–6219.

[31] Wilbourn RV, Moatt JP, Froy H, Walling CA, Nussey DH, Boonekamp JJ. The relationship between telomere length and mortality risk in non-model vertebrate systems: a meta-analysis. Phil Trans Roy Soc B: Biolog Sci. 2018;373: 20160447.

[32] Kriticos DJ, Ota N, Hutchison WD, Beddow J, Walsh T, Tay WT, et al. The potential distribution of invading *Helicoverpa armigera* in North America: is it just a matter of time?. PLoS One. 2015;10: e0119618.

[33] Pearce SL, Clarke DF, East PD, Elfekih S, Gordon KH, Jermiin LS, et al. Genomic innovations, transcriptional plasticity and gene loss underlying the evolution and divergence of two highly polyphagous and invasive *Helicoverpa* pest species. BMC Biol. 2017;15: 1–30.

[34] Feng HQ, Wu KM, Cheng DF, Guo YY. Northward migration of *Helicoverpa armigera* (Lepidoptera: Noctuidae) and other moths in early summer observed with radar in northern China. J Econ Entomol. 2004;97: 1874–1883.

[35] Arnemann JA, Roxburgh S, Walsh T, Guedes J, Gordon K, Smagghe G, et al. Multiple incursion pathways for *Helicoverpa armigera* in Brazil show its genetic diversity spreading in a connected world. Sci Rep. 2019;9: 1–2.

[36] Shadnia S, Ebadollahi-Natanzi A, Ahmadzadeh S, Karami-Mohajeri S, Pourshojaei Y, Rahimi HR. Delayed death following paraquat poisoning: three case reports and a literature review. Toxicol Res. 2018;7: 745–753.

[37] Kim J, Do Shin S, Jeong S, Suh GJ, Kwak YH. Effect of prohibiting the use of Paraquat on pesticide-associated mortality. BMC Pub Health. 2017;17: 858.

[38] Ludlow AT, Spangenburg EE, Chin ER, Cheng WH, Roth SM. Telomeres shorten in response to oxidative stress in mouse skeletal muscle fibers. J Gerontol A: Biomed Sci Med Sci. 2014;69: 821–830.

[39] Krůček T, Korandová M, Šerý M, Frydrychová RČ, Krůček T, Korandová M, et al. Effect of low doses of herbicide paraquat on antioxidant defense in *Drosophila*. Arch Insect Biochem Physiol. 2015;88: 235–248.

[40] Korandová M, Krůček T, Szakosová K, Kodrík D, Kühnlein RP, Tomášková J, et al. Chronic low-dose pro-oxidant treatment stimulates transcriptional activity of telomeric retroelements and increases telomere length in *Drosophila*. J Insect Phys. 2018;104: 1–8.

[41] Zhang XS, Wang T, Lin XW, Denlinger DL, Xu WH. Reactive oxygen species extend insect life span using components of the insulin-signaling pathway. Proc Natl Acad Sci. 2017;114: E7832–E7840.

[42] Apirajkamol N, James B, Gordon KH, Walsh TK, McGaughran A. Oxidative stress delays development and alters gene expression in the agricultural pest moth, *Helicoverpa armigera*. Ecol Evol. 2020;10: 5680–5693.

[43] Okazaki SA, Tsuchida K, Maekawa H, Ishikawa H, Fujiwara H. Identification of a pentanucleotide telomeric sequence,(TTAGG) n, in the silkworm *Bombyx mori* and in other insects. Mol Cell Biol. 1993;13: 1424–1432.

[44] Frydrychová R, Grossmann P, Trubac P, Vítková M, Marec FE. Phylogenetic distribution of TTAGG telomeric repeats in insects. Genome. 2004;47: 163–178.

[45] Sahara K, Marec F, Traut W. TTAGG telomeric repeats in chromosomes of some insects and other arthropods. Chromosome Res. 1999;7: 449–460.

[46] Livak KJ, Schmittgen TD. Analysis of relative gene expression data using real-time quantitative PCR and the 2−ΔΔCT method. Methods. 2001;25: 402–408.

[47] Schmittgen TD, Livak KJ. Analyzing real-time PCR data by the comparative C T method. Nature Prot. 2008;3: 1101.

[48] Nersisyan L, Arakelyan A. Computel: computation of mean telomere length from whole-genome next-generation sequencing data. PLoS One. 2015;10: e0125201.

[49] Anderson CJ, Tay WT, McGaughran A, Gordon K, Walsh TK. Population structure and gene flow in the global pest, *Helicoverpa armigera*. Mol Ecol. 2016;25: 5296–5311.

[50] Corp IB. IBM SPSS statistics for windows, version 22.0. Armonk, NY: IBM Corp. 2013.

[51] Wickham H. ggplot2: elegant graphics for data analysis. Springer; 2016.

[52] R Core Team. R: A language and environment for statistical computing. 2019.

[53] Jemielity S, Kimura M, Parker KM, Parker JD, Cao X, Aviv A, et al. Short telomeres in short-lived males: what are the molecular and evolutionary causes?. Aging Cell. 2007;6: 225–233.

[54] Herborn KA, Heidinger BJ, Boner W, Noguera JC, Adam A, Daunt F, et al. Stress exposure in early post-natal life reduces telomere length: an experimental demonstration in a long-lived seabird. Proc Roy Soc B: Biolog Sci. 2014;281: 20133151.

[55] Simon NM, Smoller JW, McNamara KL, Maser RS, Zalta AK, Pollack MH, et al. Telomere shortening and mood disorders: preliminary support for a chronic stress model of accelerated aging. Biolog Psych. 2006;60: 432–435.

[56] Kiecolt-Glaser JK, Gouin JP, Weng NP, Malarkey WB, Beversdorf DQ, Glaser R. Childhood adversity heightens the impact of later-life caregiving stress on telomere length and inflammation. Psychosomatic Med. 2011;73: 16.

[57] Entringer S, Epel ES, Lin J, Buss C, Shahbaba B, Blackburn EH, et al. Maternal psychosocial stress during pregnancy is associated with newborn leukocyte telomere length. Am J Ob Gynecol. 2013;208: 134–e1.

[58] Kotrschal A, Ilmonen P, Penn DJ. Stress impacts telomere dynamics. Biol Lett. 2007;3: 128–130.

[59] Geiger S, Le Vaillant M, Lebard T, Reichert S, Stier A, Le Maho Y, et al. Catching-up but telomere loss: half-opening the black box of growth and ageing trade-off in wild king penguin chicks. Mol Ecol. 2012;21:1500–1510.

[60] Aydinonat D, Penn DJ, Smith S, Moodley Y, Hoelzl F, Knauer F, et al. Social isolation shortens telomeres in African Grey parrots (*Psittacus erithacus erithacus*). PLoS One. 2014;9: e93839.

[61] Boonekamp JJ, Mulder GA, Salomons HM, Dijkstra C, Verhulst S. Nestling telomere shortening, but not telomere length, reflects developmental stress and predicts survival in wild birds. Proc Roy Soc B: Biolog Sci. 2014;281: 20133287.

[62] Parolini M, Romano A, Costanzo A, Khoriauli L, Santagostino M, Nergadze SG, et al. Telomere length is reflected by plumage coloration and predicts seasonal reproductive success in the barn swallow. Mol Ecol. 2017;26: 6100–6109.

[63] Young RC, Welcker J, Barger CP, Hatch SA, Merkling T, Kitaiskaia EV, et al. Effects of developmental conditions on growth, stress and telomeres in black-legged kittiwake chicks. Mol Ecol. 2017;26: 3572–3584.

[64] Rollings N, Miller E, Olsson M. Telomeric attrition with age and temperature in Eastern mosquitofish (*Gambusia holbrooki*). Naturwissenschaften. 2014;101: 241–244.

[65] Simide R, Angelier F, Gaillard S, Stier A. Age and heat stress as determinants of telomere length in a long-lived fish, the Siberian sturgeon. Physiolog Biochem Zool. 2016;89: 441–447.

[66] Zhang Q, Han X, Hao X, Ma L, Li S, Wang Y, et al. A simulated heat wave shortens the telomere length and lifespan of a desert lizard. J Therm Biol. 2018;72: 94–100.

[67] Burraco P, Díaz-Paniagua C, Gomez-Mestre I. Different effects of accelerated development and enhanced growth on oxidative stress and telomere shortening in amphibian larvae. Sci Rep. 2017;7: 1–1.

[68] Lee JR, Xie X, Yang K, Zhang J, Lee SY, Shippen DE. Dynamic interactions of *Arabidopsis* TEN1: stabilizing telomeres in response to heat stress. Plant Cell. 2016;28: 2212–2224.

[69] Romano GH, Harari Y, Yehuda T, Podhorzer A, Rubinstein L, Shamir R, et al. Environmental stresses disrupt telomere length homeostasis. PLoS Genet. 2013;9: e1003721.

[70] Beletsky AV, Malyavko AN, Sukhanova MV, Mardanova ES, Zvereva ME, Mardanov AV, et al. Expression of genes involved in DNA repair and telomere maintenance in the yeast *Hansenula polymorpha* DL1 under heat stress. Doklady Biochem Biophys. 2015;462: 185–188.

[71] Noreikiene K, Kuparinen A, Merilä J. Age at maturation has sex-and temperature-specific effects on telomere length in a fish. Oecologia. 2017;184: 767–777.

[72] Debes PV, Visse M, Panda B, Ilmonen P, Vasemägi A. Is telomere length a molecular marker of past thermal stress in wild fish?. Mol Ecol. 2016;25: 5412–5424.

[73] Louzon M, Pauget B, Gimbert F, Morin-Crini N, de Vaufleury A. Ex situ environmental risk assessment of polluted soils using threshold guide values for the land snail *Cantareus aspersus*. Sci Total Environ. 2020;721: 137789.

[74] Arimura GI, Kost C, Boland W. Herbivore-induced, indirect plant defences. Biochim Biophys Acta Mol Cell Biol Lipids. 2005;1734: 91–111.

[75] Mason JM, Reddy HM, Frydrychova RC. Telomere maintenance in organisms without telomerase. In: Seligmann H, editor. DNA replication—current advances. Rijeka, Croatia: InTech; 2011. pp. 323–346.

